# Epigenetic mechanism of carbohydrate sulfotransferase 3 (*CHST3*) downregulation in the aging brain

**DOI:** 10.1101/741355

**Authors:** David Baidoe-Ansah, M Sadman Sakib, Shaobo Jia, Andre Fischer, Rahul Kaushik, Alexander Dityatev

**Author notes:** Corresponding authors: Dr. Rahul Kaushik, Dr. Alexander Dityatev, German Center for Neurodegenerative Diseases (DZNE), Leipziger Str. 44, Haus 64, 39120 Magdeburg, Germany.

## Abstract

Neural extracellular matrix (ECM) is a complex molecular meshwork surrounding neurons and glial cells in the extracellular space. Structural and functional state of ECM in the brain is tightly regulated by various components of neural ECM such as hyaluronic acid, chondroitin sulfate proteoglycans, link proteins, tenascins, various matrix-modifying enzymes such as chondroitin sulfate synthases and carbohydrate sulfotransferase together with matrix-degrading enzymes. Age-dependent accumulation of ECM molecules is implicated in the age-associated decline in synaptic and cognitive functions. Understanding age-associated changes in the expression of genes involved in regulating various components of ECM can provide an insight into the role of ECM in the aging brain. Hence, in this study, we compared the expression levels of ECM regulating genes in three groups of mice: 2-3 months old mice (2-3M), 22- to 26-month-old mice (22-26M) and more than 30-month-old mice (>30M). Using qPCR, we discovered that in the hippocampus of >30M old mice, the majority of ECM related genes are downregulated, while genes related to neuroinflammation are highly upregulated. This pattern was accompanied by a decrease in cognitive performance of the >30M old mice and was most correlated among ECM-related genes with the downregulation of carbohydrate sulfotransferase 3 (*CHST3*) gene expression. Interestingly, in 24-26M mice, no general decrease in the expression of ECM related genes was observed, although we still found the upregulation in neuroinflammatory genes and downregulation of *CHST3*. Further analysis of epigenetic mechanisms revealed a decrease in H3K4me3, three methyl groups at the lysine 4 on the histone H3 proteins, associated with the promoter region of *CHST3* gene in non-neuronal (NeuN-negative) but not in neuronal (NeuN-positive) cells. We conclude that in 22-26 M old brains there are minor changes in expression of the studied *bona fide* neural ECM genes but there is a prominent epigenetic dysregulation of the *CHST3* gene responsible for 6-sulfation of chondroitin sulfates, which may lead to impaired brain plasticity and cognitive decline.

## Introduction

Plasticity in the nervous system underlies the ability of the brain to learn and modify the existing behavior and allow the organism to adapt to the changing environments (Burke & Barnes, 2006; Sengpiel, 2007; Galván, 2010; Attardo *et al.*, 2018; Ribic *et al.*, 2019). However, this ability of the brain decreases with age and highly correlates with the age-related gradual increase in the accumulation of the brain ECM (Dityatev & Fellin, 2008; Vegh *et al.*, 2014). ECM in the brain exists in a condensed form known as perineuronal nets (PNN), mostly present around parvalbumin (PV)-positive interneurons, and as a more diffuse perisynaptic ECM around synapses on excitatory cells (Dityatev & Schachner, 2003). Expression of both forms of ECM, i.e. perineuronal and perisynaptic, is increased significantly with age in many brain areas including both hippocampus and cortex (Dityatev *et al.*, 2007; Vegh *et al.*, 2014). During postnatal development, the formation of perineuronal nets underlies the closure of the critical period that is marked by the significant reduction in the ability of the brain to modify the already formed neuronal connections (Dityatev & Fellin, 2008; Miyata *et al.*, 2012; Carulli *et al.*, 2016). This helps to mature the brain circuitries and plays a critical role in brain function. Following this important developmental event, there is a life-long gradual accumulation of ECM proteoglycans in the brain that might play an important role in the age-dependent decline in cognitive functions such as learning and memory, which is a shared feature of both normal and pathological aging (Richard & Lu, 2019). However, the molecular mechanism behind this age-dependent increase in ECM has been elusive so far.

The backbone of neural ECM is a glycan, hyaluronic acid (HA), that is synthesized and anchored to the cell surface by hyaluronan synthases (*HAS1-4*) (Weigel, 2015). Other major constituents are chondroitin sulfate proteoglycans (CSPGs) of the lectican family, such as aggrecan (*ACAN*), brevican (*BCAN*), neurocan *(NCAN)*, and versican *(VCAN),* as well as phosphacan *(PCAN)*. CSPGs bind to HA via the amino-terminal hyaluronan binding domain, while the carboxy-terminal binding domain binds to the ECM glycoprotein tenascin-R *(TNR),* resulting in a net-like structure. Hyaluronan and proteoglycan link proteins (*HAPLN1-4*) stabilize the binding of lecticans to HA (Dityatev & Schachner, 2003).

Lecticans are proteoglycans consisting of core proteins and a variable number of glycosaminoglycan (CS-GAGs) side chains that are long, unbranched polysaccharides made of repeating disaccharide units of amino sugar, either N-acetylglucosamine (GlcNAc) or N-acetylgalactosamine (GalNAc), and glucuronic acid (GlcA). Chondroitin polymerizing factor 2 (*CHPF2*), chondroitin sulfate synthases 1 (*CHSY1*), chondroitin sulfate synthase 3 (*CHSY3*) regulate the length of glycosaminoglycans (GAG) side chains that are added to the core proteins of CSPGs. GAG chains are further matured by other highly controlled modifications such as sulfation. Various carbohydrate sulfotransferase genes such as *CHST3*, *CHST7*, *CHST11*, *CHST13* catalyze the sulfation of GAG chains mostly at position 4 (C4S) or 6 (C6S). *CHST3* and *CHST7* control the C6S whereas *CHST11* and *CHST13* regulate the C4S of the GAG chains (Mikami & Kitagawa, 2013; Miyata & Kitagawa, 2017).

The inhibitory role of ECM in age-dependent decline in brain plasticity is largely attributed to the increasing ratio of C4S/C6S of the GAG chains attached to CSPGs. Four-sulfated GAG chains seem to be highly inhibitory to axonal growth (Smith-Thomas *et al.*, 1995), whereas C6-sulfated GAG chains seem to promote axonal growth (Wang *et al.*, 2008). Hence the ratio of C4S/C6S seems to be a critical determinant of the functional implication of GAG chains on neurite outgrowth, structural plasticity and cognition and seems to be gradually increasing not only at the end of critical period but also throughout life (Carulli *et al.*, 2010; Miyata *et al.*, 2012; Foscarin *et al.*, 2017). However, the mechanisms behind the age-dependent increase in the C4S/C6S ratio are not yet clear.

Several studies have shown that there is a gradual age-dependent physiological increase in the neuroinflammation, which is related to activation of astrocytes and microglia (Godbout *et al.*, 2005; Lynch, 2010), and is also thought to be involved in cognitive decline (Pluvinage *et al.*, 2019). Such neuroinflammation is marked by increased expression of astrocytic and microglial markers, such as glial fibrillary acidic protein (GFAP) (Lyons *et al.*, 2009; George & Geller, 2018) and ionized calcium-binding adaptor molecule 1 (IBA1) respectively (Ito *et al.*, 2001) along with the upregulation of proinflammatory cytokines such as interleukin 6 (IL-6) and tumor necrosis factor alpha (TNFα) (Lynch, 2010). Additionally, pathological conditions such as brain injury also lead to increased activation of astrocytes that stimulates secretion of several ECM molecules that ultimately results in the formation of the glial scar (Fawcett & Asher, 1999; Beggah *et al.*, 2005; Cregg *et al.*, 2014; George & Geller, 2018). However, whether the increased activation of astrocytes under the conditions of physiological aging will also result in increased ECM production remain to be investigated.

Thus, with the current study, we aimed to investigate the age-dependent alteration in the expression of ECM and related genes, including genes that regulate the expression of GAG chains and their sulfation. Moreover, we correlated the expression of ECM related genes to cognitive decline and neuroinflammation related genes. We used qRT-PCR as a highly sensitive and reliable method to be able to detect even minor systematic age-dependent differences in gene expression. Furthermore, we also investigated the potential underlying molecular mechanisms such as epigenetic changes that might be central to such gene expression regulation.

## Materials and methods

### Animals

All animal experiments were conducted in accordance with the ethical animal research standards defined by German law and approved by the Ethical Committee on Animal Health and Care of the State of Saxony-Anhalt, Germany (license 42502-2-1343 DZNE). The present study used 55 male C57BL6/J mice in total, which included 28 young (2 to 3-month-old, hereafter referred as 2-3M), 10 mice (24 to 26-month-old, hereafter referred as 24-26M), 8 mice (22 to 23-month-old, hereafter referred as 22-23M) and 9 mice (>30-month-old, hereafter referred as >30M). Numbers of mice used in each experiment are described in the text and figure legends. All mice were transferred to the research facility from the animal breeding house and housed individually with food and water available ad libitum for at least 72 hours before experiments under a reversed 12/12 light/dark cycle (light on 9 P.M.). All behavioral experiments were performed during the dark phase of the cycle, i.e. when mice are active, under constant temperature and humidity.

### Experimental design

In the first batch of animals, we measured the cognitive decline in >30M mice compared to 2-3M mice. All mice were sacrificed and brains were used for the gene expression analysis. To further investigate the changes in the expression of ECM genes at the earlier stages of aging, we conducted the gene expression analysis on the second batch of twenty naive animals: ten 2-3M and ten 24-26M mice. Additionally, tissue samples from eight 2-3M and eight 22-23M mice were further used for FACS sorting and analysis of cell type-specific epigenetic changes of the selective genes.

### Behavioral analysis

All behavioral tests were done on the first batch of animals, which consisted of nine >30M mice and ten 2-3M mice. Anymaze 4.99 (Stoelting Co., Wood Dale, IL, USA) was used for capturing and analyzing animal’s performance during behavioral tests. Animals were subsequently tested using open field test, novel object location test (NOLT) and the novel object recognition test (NORT).

### Open field

Animals from each experimental group were put into an open field arena (50 × 50 × 30 cm) (Holter *et al.*, 2015; Kaushik *et al.*, 2018) and allowed to freely move for 10 minutes while being recorded by an overhead camera. The arena was predefined into two parts, the central area (30 × 30 cm) and a peripheral area (the 10 cm area adjacent to the wall of recording chamber). Total distance moved and time spends in the central/peripheral area were evaluated for animal’s general activity and anxiety.

### Novel object location test

The novel object location test was carried out in the same open field arena. This test includes two phases: the encoding phase and retrieval phase (Vogel-Ciernia & Wood, 2014). In the encoding phase, animals were allowed to explore the arena with a pair of identical objects for 10 minutes. 24 hours later, in the retrieval phase, the animals were given 10 minutes to explore the arena again with the same objects, except that one of them was placed in a novel position. Exploring time for an object located at familiar (F) and novel (N) position as well as discrimination ratio [(N-F)/(N+F)] × 100 % were used to analyze animal’s behavior.

### Novel object recognition task

The novel object recognition test was also conducted in the open field arena containing the encoding phase and retrieval phase as described previously (Antunes & Biala, 2012; Kaushik *et al.*, 2018). Briefly, in the encoding phase, animals were allowed to explore the arena with a pair of identical objects. 24 hours later, in the retrieval phase, one of the object was replaced by a novel object that the animal hadn’t seen before. Both the encoding and retrieval phase lasted 10 minutes. Exploration time in seconds for both objects was further analyzed by a trained observer who was blind to treatment conditions. Exploring time for familiar (F) and novel (N) objects, as well as discrimination ratio [(N-F)/(N+F)] × 100 %, were used to evaluate animals’ recognition memory.

### Tissue isolation

Animals were quickly decapitated and brain tissue was isolated in ice-cold PBS. Specific brain regions were further dissected and snap-frozen on dry ice and stored in −80°C until further use or fixed. Hippocampal and cortical samples were isolated and processed as described earlier. Left and right hippocampi from the second batch of animals were stored for only RNA extraction and downstream processing. For symmetry, we used the right hippocampus and right cortex for subsequent analysis.

### RNA extraction, cDNA conversion, and QPCR

Total RNA was extracted from the frozen brain regions using EURx GeneMatrix DNA/RNA Extracol kit (Roboklon Cat. No. E3750) according to the manufacturer’s recommendations (Ventura Ferreira *et al.*, 2018). The RNA yield, purity, and integrity were determined with Nano-drop and gel electrophoresis respectively to confirm the absence of genomic DNA. Furthermore, 1.5 µg of RNA was used for cDNA conversion using the High-Capacity cDNA Reverse Transcription Kit (Cat.4368814). RT-qPCR was carried out using TaqMan gene expression array (Cat. 4331182) from Thermo-Fisher Scientific using Quant-Studio-5 from Applied Biosystems. Nineteen genes were analyzed, comprising five CSPG’s genes, namely *ACAN*, *BCAN*, *NCAN, VCAN and PCAN*, two genes coding for link proteins *HAPLN1* and *TNR*, eight genes regulating the synthesis of HA, GAG’s and their sulfation levels such as *HAS2*, *CHPF2*, *CHSY1*, *CHSY3*, *CHST3*, *CHST7*, *CHST11,* and *CHST13,* respectively. Additionally, three cell type-specific markers such as aldehyde dehydrogenase 1 family member L1 (*ALDH1L1*), glial fibrillary acidic protein (*GFAP*), ionized calcium-binding adaptor molecule 1 (*IBA1*) and two inflammatory markers; interleukin 6 (*IL6*) and tumor necrosis factor (*TNF*) were also quantitatively measured and analyzed relative to expression of Glyceraldehyde 3-phosphate dehydrogenase (*GAPDH*) (Meldgaard *et al.*, 2006). The mRNA levels of *GAPDH* were not significantly changed between 2-3M, 24-26M and >30M ice making it a suitable candidate for the normalization of other genes. Additionally, due to the use of some cortical samples for optimization of RNA extraction protocol and insufficient cDNA for some of the genes, we could only run 5 samples per group for both *BCAN* and *HAPLN1*. As such, we could not determine the mRNA levels of *PCAN* in 2-3M and 24-26M groups.

### FACS sorting for neuronal and glial nuclei extraction

Hippocampi from 2-3M and 22-23M mice from the third batch of animals were dissected, flash frozen and kept at −80°C until further use. The nuclei extraction protocol was adapted from (Halder *et al.*, 2016) with slight modifications. Briefly, left hippocampi were used for the extraction of nuclei in order to perform chromatin immunoprecipitation (ChIP). Frozen tissues were homogenized in low-sucrose buffer (LSB; 5mM CaCl2 -Applichem A3652,0500, 320mM sucrose-Applichem A4737,5000, 0.1mM EDTA-Invitrogen AM9260G, 5mM MgAc2 -Sigma M5661-250G, 10mM HEPES pH 8 - Gibco 15630-056, 1mM DTT – Roth 6908.2, 0.1% Triton X-100 – Sigma T8787, 1x EDTA free Roche protease inhibitor cocktail - Sigma 5056489001) in 1.5mL tubes. The crosslinking step was performed by adding 1% formaldehyde as final concentration and incubated for 10 minutes at room temperature. Furthermore, cross-linking was stopped by adding 125mM glycine (Applichem A1067,0500) with incubation for 5 minutes at room temperature. Nuclei were spun down by centrifugation at 2000 × g for 3 minutes at 4°C. The remaining crude nuclear pellet was re-suspended into LSB and homogenized with a mechanical homogenizer (IKA Ultraturax T10). The suspension was layered on 6mL high-sucrose buffer (HSB; 3mM MgAc2, 1mM DTT, 1000mM Sucrose, 10mM HEPES pH 8, protease inhibitor) in oak-ridge tubes and centrifuged for 10 minutes at 3220 × g at 4°C. After removing the upper phase containing myelin debris, the resulting nuclear pellet was re-suspended into the leftover buffer, transferred into microfuge tubes (Eppendorf DNA-low bind, 022431021) and centrifuged at 2000 × g for 3 minutes to recover the nuclear pellet. Before staining, the nuclei pellet was resuspended into PBS containing Tween-20 and BSA (PBTB; 1% BSA, 0.2% Tween-20, EDTA-free protease inhibitor dissolved in 1X PBS) and Anti-NeuN-Alexa488 conjugated antibody (MAB377X) was added at 1:1000 dilution. After 1-hour incubation at 4°C, samples were washed twice with PBS and proceeded with the sorting in BD FACS Aria III. Sorted nuclei were collected into Falcon tubes, pelleted with brief centrifugation and pellets were flash-frozen and saved at −80°C until further processing for ChIP. In parallel, right hippocampi were used to perform nuclei extraction followed by the RNA extraction using tissue from the same animals. A similar protocol was followed as described above, except the RNAse inhibitor (Promega N2615) was used in all buffers and no crosslinking was done. After sorting, RNA was extracted using Trizol reagent (Sigma T9424) and RNA was kept at −80°C until further processing.

### Chromatin immunoprecipitation and qPCR

Chromatin immunoprecipitation was performed as described previously (Halder *et al.*, 2016) with slight modifications. Briefly, 0.2µg chromatin was used for doing H3K4me3 ChIP (1µg antibody, ab8580). 1:1 mixture of Protein A and G beads were used for pre-clearing and immunoprecipitation. ChIP DNA was subjected to qPCR using SYBR green reagents with primers targeting *CHST3* transcription start site +/− 200bp. Primers were designed using primer BLAST(NCBI) by taking the genomic sequence and selecting for amplicon between 70-150bp. The primer pair used for ChIP-qPCR was (5’-3’ direction); forward: GGGCCTTTGTTCCCGACTTA and reverse: CTCGTCCTCAAGGGTAGGGA. Percent input method (% Input) was used to calculate H3K4me3 enrichment.

### cDNA preparation from sorted nuclear RNA and qPCR

RNA concentration from sorted nuclei was measured in Qubit 2.0 (Invitrogen) using RNA high sensitivity kit. Three ng of RNA was used to prepare cDNA using SMART-Seq v4 Ultra Low Input Kit (Takara). Following cDNA preparation, equal amounts of cDNA were subjected to qPCR using primers (designed by Roche universal probe library tool) for *CHST3* mRNA. Primer sequences were (5’-3’ direction): CCACAGCAGCCAGATCTTC (forward), and TGGGGGACACTCTGATCCT (reverse). A standard curve was generated to calculate primer efficiency, which was used to obtain normalized expression value (delta-delta Ct method), using DNA topoisomerase I *(TOP1)* as housekeeping gene (Penna *et al.*, 2011). Primer sequences were (5’-3’ direction): TGCCTCCATCACACTACAGC (forward), and CGCTGGTACATTCTCATCAGG (reverse:). Relative expression was calculated by normalizing to the 2-3M samples.

### Statistical analysis

Statistical analysis was performed using GraphPad Prism 7.0 (GraphPad Software Inc., La Jolla, USA) and Statistica 8.0 (StatSoft, USA) software. All data are shown as mean ± SEM with n being the number of samples (mice). Asterisks in figures indicate statistical significance (with details in the figure legend or results). The hypothesis that experimental distributions follow the Gaussian law was verified using Kolmogorov-Smirnov, Shapiro-Wilk, or D’Agostinio tests. For pairwise comparisons, we performed Students’ t-test where the samples qualify for normality test; otherwise, the Mann-Whitney test was employed. Additionally, Wilcoxon matched-pairs test was used for paired t-test that did not pass the normality test. Holm-Sidak’s multiple comparisons t-test used for independent comparisons. Spearman correlation coefficients were computed to estimate the correlation between the variables studied. The p-values represent the level of significance as indicated in figures by asterisks (*p < 0.05, **p < 0.01, ***p < 0.001 and **** p < 0.0001) unless stated otherwise.

## Results

To understand the interplay between cognitive decline, neuroinflammation and high levels of ECM proteins in aging, we investigated cognitive functions, mRNA expression of ECM molecules as well as inflammatory markers in >30M mice as compared to 2-3M mice. The mice were subsequently sacrificed to isolate hippocampus and cortex for mRNA analysis of genes related to various components of neural ECM such as hyaluronic acid, chondroitin sulfate proteoglycans, link proteins, tenascins, matrix-modifying enzymes such as chondroitin sulfate synthases, carbohydrate sulfotransferase, and matrix-degrading enzymes together with various inflammatory markers using RT-qPCR.

### Cognitive functions are impaired in >30M mice

To identify the potential interaction between the aforementioned major age-dependent phenotypes, we used NOLT and NORT tests to access the cognitive status of nine >30M and ten 2-3M mice. >30M mice covered less distance compared to 2-3M (traveling mean distance of 32.1 ± 2.3 m vs 50.6 ± 1.8 m, p < 0.001, unpaired Welch’s t-test; Figure 1B) in the open-field task. There was no difference in time spent in the central area (92.0 ± 16.7 s vs.124.7 ± 7.1 s; p = 0.1499, Mann-Whitney test; Figure 1B), suggesting no major changes in anxiety in >30M mice.

**Figure 1.**
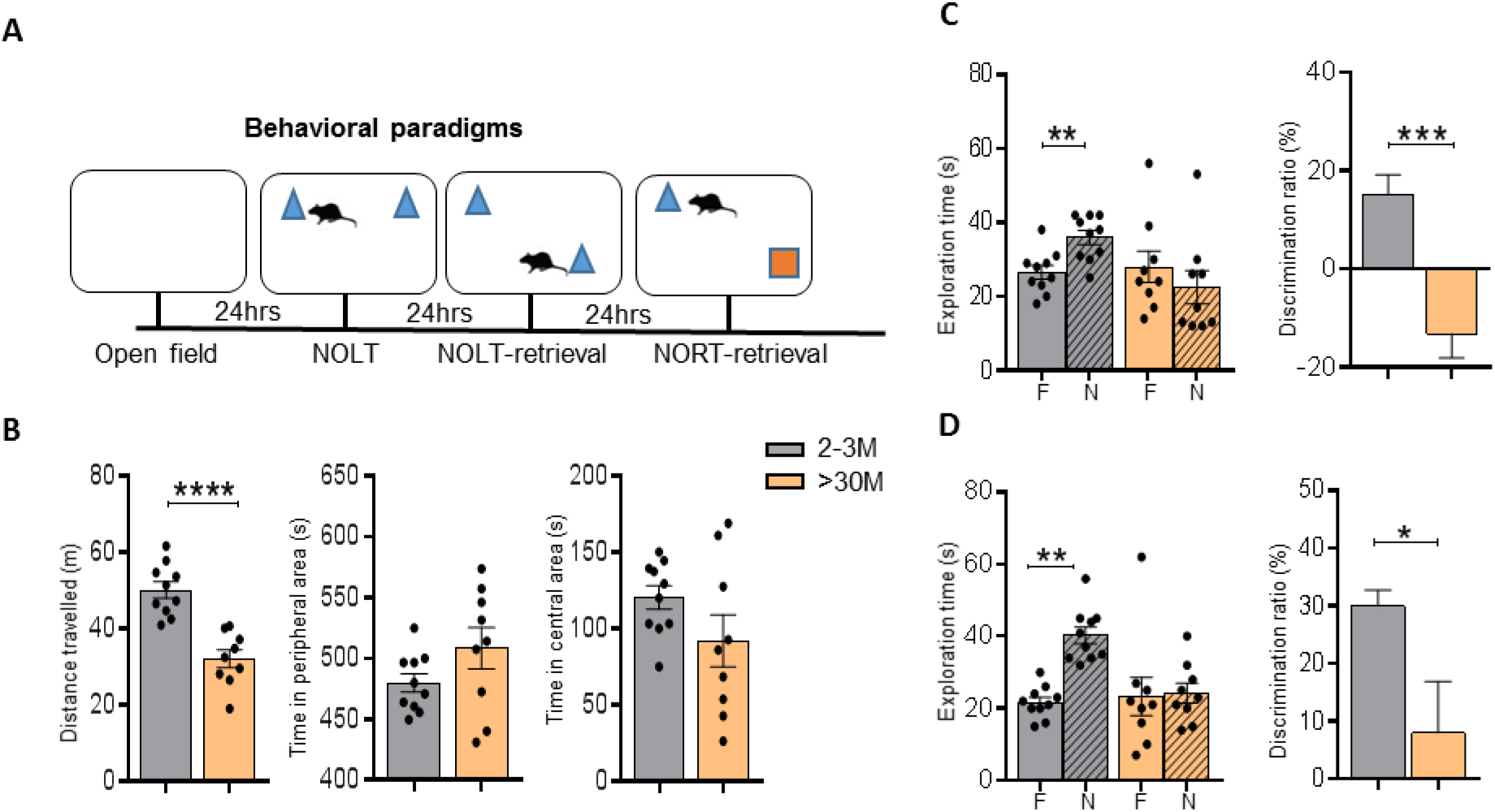
Impaired long-term memory in >30M mice. Long-term memory was tested in >30M (>30-month-old) and 2-3M (2- to 3-month-old) mice using the novel object location task (NOLT) and novel object recognition task (NORT). **(A)** Time-line for all cognitive tests. **(B)** >30M mice traveled shorter distances compared to 2-3M mice in the open field. Moreover, >30M mice also spent equal time exploring familiar (F) and novel (N) locations and objects, indicating a failure to discriminate between objects in NOLT **(C)** and NORT **(D).** The difference in discrimination between >30M and 2-3M mice was further confirmed using discrimination ratio analysis. Bar graphs show mean ± SEM values for each animal. *p* < 0.05, \*\**p* < 0.01, \*\*\**p* < 0.001 and **** *p* < 0.0001 represent significant differences between 2-3M (N=10) and >30M (N=9) mice.

In NORT, >30M mice seem to have difficulty in discriminating between the familiar (F) and novel (N) objects with exploration time 28.0 ± 4.262 s vs. 22.44 ± 4.476 s, respectively (p = 0.0508, Wilcoxon matched-pairs test). In contrast, 2-3M mice showed strong discrimination between these objects (26.5 ± 1.833 s vs. 35.9 ± 1.906 s; p = 0.0098, Wilcoxon matched-pairs test). Comparison of discrimination ratio revealed a significant difference between the 2-3M and >30M mice to distinguish novel objects (15.22 ± 3.949 vs. −13.17 ± 4.965; p = 0.0003, Mann-Whitney test; Figure 1C).

Similar to the NOLT, 2-3M mice spent a significant amount of time exploring the novel objects with exploration time of 40.4 ± 2.1 s (N) and 21.3 ± 1.3 s (F) (P = 0.0020, Wilcoxon matched-pairs test) in the NORT test. However, >30M mice spent nearly equal time exploring both objects (24.2 ± 2.7 s (N) vs. 23.3 ± 5.3 s (F); p=0.4805, Wilcoxon matched-pairs test). In line with this finding, the discrimination ratio analysis revealed a difference between >30M and 2-3M mice (30.0 ± 2.7 vs. 8.0 ± 8.9; p = 0.0182, Mann-Whitney test; Figure.1D). These data suggest a clear cognitive impairment in >30M mice in both behavioral paradigms as compared to the 2-3M animals.

### Expression of CSPGs core-proteins and enzymes regulating synthesis and sulfation of GAG chains in the hippocampus of >30M mice

CSPG core proteins are highly upregulated in the aged brain (Vegh *et al.*, 2014). We investigated if the upregulation of their mRNA transcripts might be the underlying mechanism of these high protein levels. Surprisingly, we observed a significantly low level of mRNA of the core-proteins of all CSPG genes; *ACAN, BCAN, NCAN, VCAN, PCAN* in >30M mice (p = 0.0155, 0.0434, 0.0155, 0.0281, and 0.0155 respectively; Holm-Sidak’s multiple comparisons t-test). Furthermore, the link protein gene *HAPLN1* was significantly reduced (p = 0.0081, Holm-Sidak’s multiple comparisons t-test). However, we could not observe any changes in the expression of ECM glycoprotein *TNR* and the gene encoding for the hyaluronan synthesis (*HAS2*) (p = 0.5720 and 0.1799 respectively; Holm-Sidak’s multiple comparisons t-test) (Figure. 2D). *GAPDH* gene was used to normalize the expression levels of these genes as a housekeeping gene whose expression was not altered in compared conditions.

**Figure 2.**
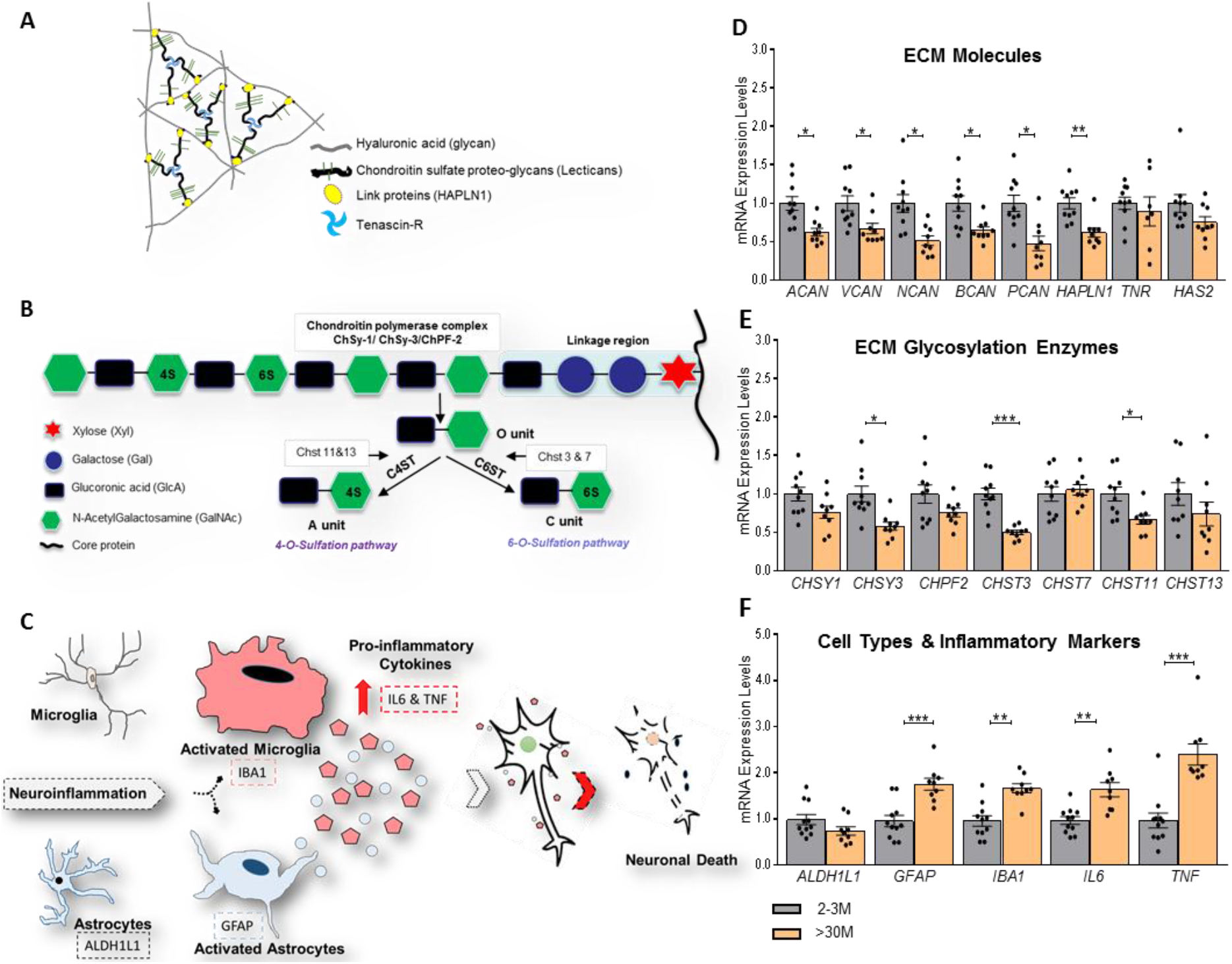
Downregulated ECM mRNA in the right hippocampus of >30M mice. **(A)** Scheme depicting the basic principle of neural ECM organization. **(B)** GAG chains attached to core proteins of various types of CSPGs (e.g. aggrecan) are polymerized by the chondroitin polymerase complex which includes the chondroitin synthase 1&3 (Chsy1&3), and chondroitin polymerizing factor 2 (Chpf2). Un-sulfated disaccharides (O unit) of GAG chains are further modified with sulfates yielding C4S (A unit) and C6S (C unit) by chondroitin 4 sulfotransferase (C4ST) and chondroitin 6 sulfotransferase (C6ST), respectively. **(C)** Neuroinflammatory microenvironment. **(D)** Genes encoding for core proteins of CSPGs; *ACAN, VCAN, NCAN, BCAN,* and *PCAN*, were down-regulated in >30M mice compared to 2-3M mice. **(E)** Moreover, in >30M mice, glycosylation enzymes were analyzed and C6 sulfotransferase *CHST3* was found to be downregulated compared to 2-3M mice. **(F)** Inflammatory cytokines and markers for glial activation were upregulated in >30M mice, a characteristic feature of aging, compared to 2-3M mice. Bar graphs show mean ± SEM values. \**p* < 0.05, \*\**p* < 0.01, \*\*\**p* < 0.001 and **** *p* < 0.0001 represent significant differences between 2-3M (N=10) and >30M (N=9) mice.

Furthermore, in order to understand the regulation of glycosylation levels of CSPGs, we investigated the expression levels of genes encoding for the glycosylating enzymes such as *CHSY1*, *CHSY3,* and *CHPF2*. Interestingly, we also observed a significant reduction in the levels of *CHSY3* (p = 0.0150, Holm-Sidak’s multiple comparisons t-test). However, we could not detect any changes in other glycosylating genes such as *CHSY1* and *CHPF2* (p = 0.2384 and 0.2612 respectively; Holm-Sidak’s multiple comparisons t-test). Moreover, in order to understand the sulfation levels of the glycoepitopes of the GAGs, we investigated the expression levels of enzymes responsible for the C6S (*CHST3*, *CHST7*) and C4S (*CHST11*, *CHST13*). Interestingly, we revealed ≈ 50% reduction in the levels of *CHST3* (p = 0.0001) *and* 34% in *CHST11* (p = 0.0381) in >30M mice (Figure 2E). In contrast, we did not see changes in *CHST7* and *CHST13* (p = 0.6485 and 0.6225 respectively; Holm-Sidak’s multiple comparisons t-test). Collectively, these data suggest an involvement of some age-associated negative feedback mechanisms that might be involved in regulating not only the expression levels of CSPGs but also the glycosylating and sulfating enzymes.

It is well established that there is an age-associated strong increase in the activation of glial cells as well as neuroinflammatory cytokines (Sochocka *et al.*, 2017). Our gene expression data also strongly support these observations, suggesting a contribution of transcriptional regulations to such upregulations. We detected a significant increase in activated astrocytic and microglia markers, *GFAP* (p = 0.0006) and *IBA1* (p = 0.0038), respectively, along with pro-inflammatory cytokines, *IL6* (p = 0.0016) and *TNF* (p = 0.0001) (Figure 2F).

### Relationships between the expression of ECM-related genes and cognitive performance

Next, we performed the correlation analysis between mRNA levels of different genes from 2-3M and >30M mice and correlated with the behavioral performance using discrimination ratio of cognitive tasks (NOLT_Discr and NORT_Discr). From the Spearman correlation matrix, we detected the strongest negative correlation between inflammatory marker genes such as GFAP, IBA1, and IL6 with NORT_Discr and NOLT_Discr with correlation coefficients −0.691, −0.818 and −0.753, respectively (Figure 3A). Also, there was a strong positive correlation between the expression of many ECM related genes. The strongest positive correlation between behavior and ECM related measures was observed between the ECM glycosylation gene *CHST3* and NOLT_Discr (r=0.619; p=0.006; Figure 3B), although not between *CHST3* and NORT_Discr (r=0.350; p=0.142; Figure 3C). These data suggest that downregulation of *CHST3* might be involved in the mechanisms associated with age-dependent cognitive decline.

**Figure 3.**
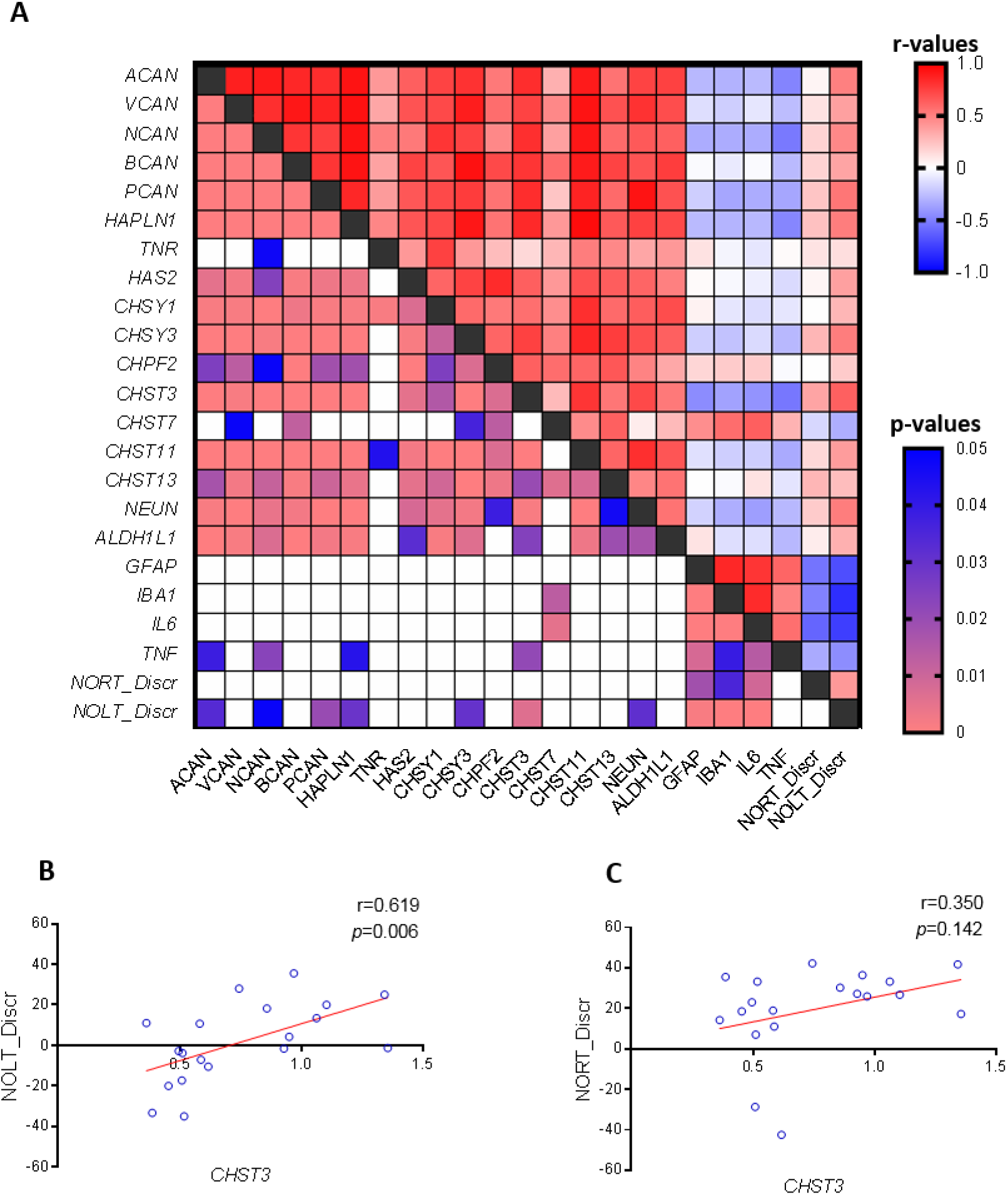
Correlation between *CHST3* and NOLT discrimination ratio. The interplay between expression of CSPGs core proteins, enzymes involved in CSPGs biosynthesis as well as sulfation, inflammatory cytokines and cognitive function (NOLT and NORT discrimination ratio) was analyzed using Spearman coefficients of correlation. **(A)** Matrix for correlation coefficients (upper half) and p-values (lower half) between all major parameters studied. The correlations are color-coded (red and blue for positive and negative correlations respectively with white for neutral correlation). **(B)** The strongest positive correlation observed between C6 sulfotransferase gene *CHST3* and NOLT_Discrimination ratio. **(C)** Correlation between C6 sulfotransferase gene *CHST3* and NORT_Discrimination ratio is not significant and shown for comparison.

### Expression of CSPGs core-proteins and enzymes regulating synthesis and sulfation of GAG chains in the hippocampus of 24-26M mice

Our gene expression data from >30M mice suggested that high levels of ECM proteins might impart a strong negative feedback mechanism in the hippocampus that regulates the expression of ECM related genes (Bonnans *et al.*, 2014). As these animals were at a very advanced stage of aging (>30 months), we hypothesize that these feedback mechanisms regulating the expression of ECM related genes might not be active at the earlier stages of aging. Hence, we investigated the gene expression analysis in 24-26M mice. Interestingly, at this age, we observed no difference in mRNA levels of core-proteins of CSPGs in the hippocampus of these mice (Figure 4A). Moreover, cortex also showed similar gene expression pattern, suggesting the brain-wide mechanisms (Figure 4B). In 24-26M mice, we observed a significant increase in the mRNA of glycosylation enzyme; *CHPF2* (p = 0.0373; Holm-Sidak’s multiple comparisons t-test) while no change in the expression of *CHSY1* and *CHSY3* (p = 0.4161 and 0.6776 respectively; Holm-Sidak’s multiple comparisons t-test). These changes were detected only in the hippocampus and not in the cortex (Figure 4C&D), suggesting some brain region-specific regulations. Remarkably, we consistently observed a strong downregulation of the mRNA levels of the *CHST3* gene in both hippocampus and cortex (p <0.0001 and p = 0.0099 respectively; Holm-Sidak’s multiple comparisons t-test). Additionally, we observed a significant upregulation of *CHST13* gene that encodes for chondroitin 4 sulfotransferase (p = 0.0325; Mann-Whitney test) only in the cortex and not in the hippocampus (Figure 4C&D), which might have an effect on system consolidation of long-term memories in the cortex.

**Figure 4.**
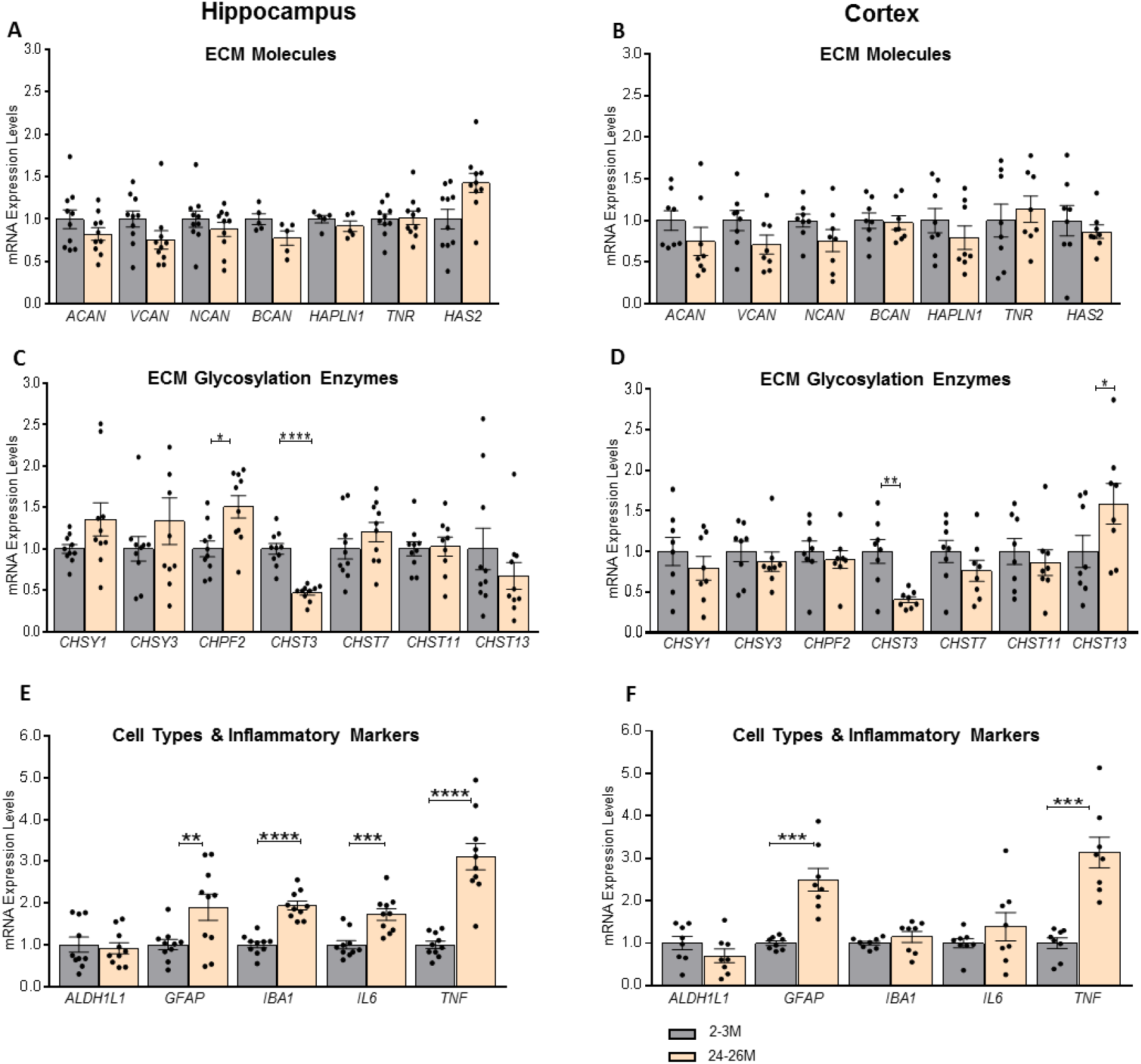
Downregulation of C6 sulfotransferase *CHST3* in hippocampus and cortex of 24-26M mice. Expression of genes involved in the biosynthesis of ECM related molecules in 24-26M mice was investigated using RT-qPCR and normalized to individual GAPDH values. **(A, B)** Genes for core-proteins of CSPGs - *ACAN, VCAN, NCAN,* and *BCAN -* as well as glycoproteins *HAPLN1*, *TNR*, and hyaluronan synthase *HAS2* were not significantly different in both hippocampus and cortex. **(C)** Genes involved in CSPGs GAG chain polymerization were slightly upregulated in the hippocampus, whereas C6 sulfotransferase *CHST3* was downregulated in the hippocampus of 24-26M mice. **(D)** However, in the cortex, polymerization complex enzymes were not significantly different, but C6 sulfotransferase *CHST3* was downregulated and C4 sulfotransferase *CHST13* was upregulated in 24-26M mice compared to 2-3M mice. **(E, F)** Inflammatory cytokines and markers for glial activation were upregulated in both hippocampus and cortex of 24-26M mice, except the cortical expression of *IBA1* and *IL6*. Bar graphs show mean ± SEM values. \**p* < 0.05, \*\**p* < 0.01, \*\*\**p* < 0.001 and **** *p* < 0.0001 represent significant differences between 2-3M (N=10 and 8) and 24-26M (N=10 and 8) mice for hippocampus and cortex, respectively.

Analysis of neuroinflammation/cell population markers revealed an increased expression of astrocytic activation marker *GFAP* in both hippocampus and cortex of these mice (p = 0.0079 and p = 0.00013, respectively; Holm-Sidak’s multiple comparisons t-test). Interestingly, we also observed an increased expression of microglial activation marker *IBA1* and inflammatory marker *IL6,* but only in hippocampus (p < 0.0001 and p = 0.0003, respectively; Holm-Sidak’s multiple comparisons t-test) and not in the cortex (p = 0.2997 and p = 0.2817 respectively; Holm-Sidak’s multiple comparisons t-test). These data suggest that aging-associated increased activation of microglia and elevation in inflammatory markers, such as *IL6,* might happen in the hippocampus before it happens in the cortex (Figure 4C&D). However, another inflammatory cytokine gene *TNF* was up-regulated in both hippocampus and cortex (p < 0.0001 and p = 0.00013 respectively; Holm-Sidak’s multiple comparisons t-test; Figure 4E&F). These data suggest that downregulation of *CHST3* might be one of the early ECM-related mechanism that is affected by aging.

### Age-dependent epigenetic changes of the CHST3 promoter in non-neuronal cells

Next, we wanted to investigate the mechanisms behind the expression of the *CHST3* gene in a cell-specific manner. Hippocampal tissue from 2-3M and 22-23M mice were sorted using fluorescence-activated cell sorting (FACS) technique with NeuN antibody conjugated to Alexa-Flour 488-A (Figure 5A) to have two population of nuclei; NeuN+ (neuronal) and NeuN− (non-neuronal). We detected a higher proportion of NeuN+ cells than NeuN− in the hippocampus of the mice brain with significantly varying proportions; including 80% (NeuN+) vs. 20% (NeuN−) in 2-3M compared to 70% (NeuN+, neurons) vs. 30% (NeuN-, glia cells) in 22-23M mice (Figure 5B). Furthermore, we performed RT-qPCR in the sorted cells to check the mRNA levels of *CHST3* expression. Interestingly, *CHST3* expression was more than 10-fold higher in glial cells when compared to neurons (Fig 5C). We detected a significant decrease in mRNA levels of *CHST3* gene in NeuN− cell population but not in NeuN+ cell population of 22-23M mice (p = 0.0200 and p = 0.3126, respectively; Figure 5C). These data indicate that the observed decrease in the expression of *CHST3* upon aging most likely originates from the glial cell population in the 22-23M mice hippocampus.

**Figure 5.**
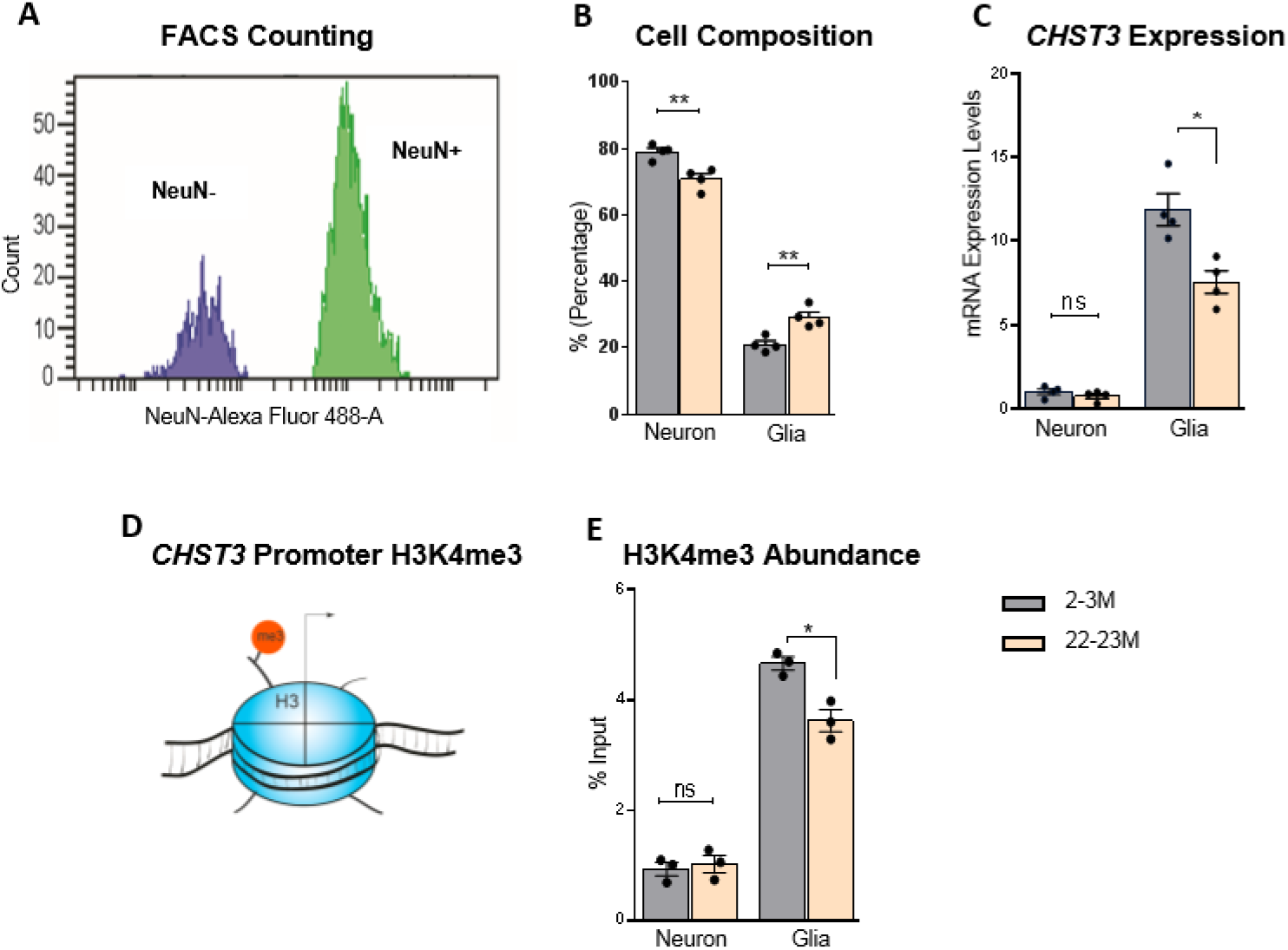
H3K4me3 is mostly enriched at the glial *CHST3* promoter in the hippocampus of 22-23M mice. Changes to *CHST3* gene expression during aging were investigated specifically at the *CHST3* promoter in NeuN positive and negative cells, which corresponds to neuronal and glial cells, respectively, in the hippocampus of 22-23M and 2-3M mice. (**A**) Representative neuronal and glial nuclei amount for each was quantified and the cell composition for each study group was determined using FACS techniques with neuron-specific NeuN-antibody conjugated to Alexa Flour 488. **(B)**Thus, percentages of neurons and non-neuronal cells (predominantly glia) within test groups were compared. (**C**) No significant difference was observed in cell type-specific nuclear RNA-qPCR for the expression of C6 sulfotransferase *CHST3* in neurons, which was, however, downregulated in glia of 22-23M mice. (**D**) Gene activation epigenetic mark, H3K4me3 at the *CHST3* promoter was investigated. (**E**) A statistical summary showing that using cell type-specific ChIP-qPCR, H3K4me3 abundance at *CHST3* promoter in neurons was found to be not different between 2-3M and 22-23M mice, whereas H3K4me3 was significantly less associated with *CHST3* promoter in glia of 22-23M mice. Bar graphs show mean ± SEM values. \**p* < 0.05, \*\**p* < 0.01, \*\*\**p* < 0.001 and **** *p* < 0.0001 represent significant differences between 2-3M (N=4 and 3 pairs) and 22-23M (N=4 and 3 pairs) mice for cell composition/*CHST3* expression and H3K4me3 abundance, respectively.

In order to understand the mechanism behind the downregulation of *CHST3*, we investigated the possible epigenetic modifications that might be involved in regulating these changes in *CHST3* expression during aging, as histone methylation has been linked to cognitive function (Kerimoglu *et al.*, 2017; Cruz *et al.*, 2018; Pu *et al.*, 2018; Mei *et al.*, 2019). Thus, we checked for the changes in the activating H3K4me3 modification at the promoter region of the *CHST3* gene in both neuronal (NeuN+) and non-neuronal (NeuN−) cells. This is a very well-characterized epigenetic modification that has been shown to be centrally involved in regulating gene expression in eukaryotes (Santos-Rosa *et al.*, 2002; Howe *et al.*, 2017). Similar to the mRNA levels of *CHST3*, levels of H3K4me3 at the promoter of *CHST3* gene in NeuN− glial sample were significantly higher in both 2-3M and 22-23M groups as compared to NeuN+ neuronal samples (Figure 5 C, E). Moreover, we observed a significant reduction in the levels of H3K4me3 on the promoter of *CHST3* specifically in NeuN− glial sample cells but not in NeuN+ neuronal samples of 22-23M mice (p = 0.6652 and p = 0.0220, respectively; unpaired t-test; Figure 5E). These data highlight specific epigenetic changes at the promoter of *CHST3* in NeuN− cell population might underlie the age-associated downregulation of *CHST3*.

## Discussion

In this study, we aimed to decipher transcriptional changes in the expression of ECM related genes that might underlie the age-dependent accumulation of neural ECM. Hence, we performed RT-qPCR to quantify the mRNA levels of different ECM molecules from three different age group of mice brain. To our surprise, we did not observe any increase in the expression of CSPG genes in the brains of the 24-26M animals. On the contrary, there seems to be a tendency for the decrease in the expression. In >30M animals, there is a significant reduction in the mRNA levels of multiple major CSPG’s genes, suggesting that an increase in brain ECM is potentially not due to increased expression of ECM genes. On the contrary, there might be a feedback mechanism suppressing the expression of these genes. Even though there is less mRNA for expression of ECM related genes upon aging, the increased levels of ECM proteoglycans suggest that the pathways regulating the degradation of ECM might be the major mechanism involved in the age-related accumulation of ECM.

Both neurons and astrocytes secrete ECM molecules in the extracellular space and contribute to the state of ECM in the brain (Dzyubenko *et al.*, 2016; Song & Dityatev, 2018). Several studies have reported that the increased activation of astrocytes leads to the secretion of large amounts of ECM molecules after brain injury (George & Geller, 2018), which ultimately result in the formation of the glial scar (Bonneh-Barkay & Wiley, 2009). However, in the aging brain, though there is increased activation of astrocytes, as indicated by elevated levels of *GFAP* expression (Clarke *et al.*, 2018), the decrease in ECM mRNA expression strongly suggests against the notion that age-dependent increase in astrocytic activation might lead to an increase in the production of ECM related genes.

A gradual increase in brain inflammation is the hallmark of the aging brain as reported previously in 24-months-old mice (Clarke *et al.*, 2018; Bok *et al.*, 2019). In accordance with the previous studies, we found an increased expression of inflammatory genes such as *GFAP*, *IBA1*, *IL6*, *TNF* in both groups of 24-26M and >30M as compared to the 2-3M animals. The increased expression of *GFAP* suggests increased activation of astrocytes. Noteworthy, we did not observe any changes in the expression of another astrocytic marker *ALDH1L1*, which is a pan-astrocyte marker, hence making it a good candidate to relate to the total number of astrocytes, which is independent of the activation state of the astrocytes (Cahoy *et al.*, 2008; Sun *et al.*, 2017). These data suggest that the number of astrocytes does not change dramatically, however; there is increased activation of astrocytes not resulting in an elevated synthesis of the studied ECM molecules.

Furthermore, we have found a significant increase in the expression levels of genes responsible for the addition of the GAG side chains such as *CHPF2* in the 24-26M animals. However, we could not observe this increase in >30M animals; rather we observed a significant decrease in the expression of *CHSY3*. These data suggest that at the initial stages of aging, the ECM molecules might be secreted with longer GAG chains. However, in the later stages of aging, there is not only the decreased expression of ECM core proteins but they also potentially carry shorter GAG chains, further suggesting the potential involvement of feedback mechanisms that might regulate not only the expression of core proteins but also the length of GAG chains associated with them. These conclusions are merely based on the expression levels of these enzymes and we cannot rule out the additional level of regulation due to the activity levels of these enzymes.

Importantly, we discovered that there is a significant and consistent decrease in the expression of *CHST3* mRNA in both 24-26M as well as the >30M brain. The chondroitin-6 sulfotransferase Chst3 is a Golgi resident enzyme that is responsible for the C6-sulfation of the GAG’s chains of CSPGs (Mikami & Kitagawa, 2013; Mizumoto *et al.*, 2014; Miyata & Kitagawa, 2017). Additionally, we found an increase in the expression of *CHST13*, which is one of the enzymes responsible for C4-sulfation of the GAGs. These data suggest that the age-dependent increased ratio of C4S/C6S might potentially be due to not only an increase in the expression of enzymes carrying out C4S but also due to a decrease in the expression of enzymes carrying out C6S. However, in >30M mice, also the expression levels of *CHST13* are unchanged. Additionally, we see a decrease in the expression of *CHST3* and *CHST11,* potentially due to an age-dependent feedback mechanism.

Collectively, our data suggest that as aging progresses the secreted CSPGs not only might have less C6-sulfated GAG chains but also shorter GAGs due to lower expression of GAG-adding enzyme complex of *CHSY3*, *CHSY1*, and *CHPF2* (Bowman & Bertozzi, 1999; Li *et al.*, 2001). Several studies have shown the central role of ECM, especially in the form of the PNNs, in the regulation of age-dependent cognitive decline, a reversal of fear memories, ocular dominant plasticity and cognitive flexibility (Dityatev *et al.*, 2010; Happel & Frischknecht, 2016; Reichelt *et al.*, 2019). However, most of the available studies are based on digestion of ECM with chondroitinase ABC (chABC)/hyaluronidase that leads to digestion of inhibitory GAG’s from both PNN as well as perisynaptic ECM. The increased ratio of C4S/C6S on the GAG’s that are present on PNN forming core protein aggrecan attracted the most attention as a potential mechanism behind the plasticity-restraining function of ECM. Moreover, the increase in the ratio is largely attributed to the decrease in C6S GAG’s (Foscarin *et al.*, 2017; Pudelko *et al.*, 2019). Here, we demonstrate that *CHST3* expression is strongly downregulated both in the cortex and the hippocampus of 24-26M mouse brain compared to the 2-3M brain. Importantly, we observed a decreased expression of *CHST3* mRNA specifically in the non-neuronal (glial) cell types along with the decrease of the expression-promoting epigenetic marker. We suggest that age-dependent decrease in C6S might be the result of a decrease in *CHST3* expression especially in non-neuronal cells of the brain.

As a first step to dissect the mechanisms by which upregulation of ECM affects cognitive performance in aging, we investigated the cognitive decline in >30M mice and correlated their performance with the expression levels of various ECM related genes. Strikingly, the highest correlation among the ECM-related genes was found for *CHST3*. In line with our findings, studies using transgenic mice overexpressing *CHST3* observed a reduction in C4S/C6S ratio. This resulted in an impairment of PNN formation and reduced maturation of PV interneurons, including a reduction in their inhibitory effects (Miyata & Kitagawa, 2016). Additionally, *CHST3* overexpression resulted in persistent cortical plasticity (Miyata *et al.*, 2012), which is contrary to downregulation of *CHST3* during early development, which leads to the closure of critical period and suppression of developmental plasticity. Moreover, we consistently found that *CHST3* gene seems to be highly downregulated not only in the hippocampus and cortex of 24-26M mice but also in the hippocampus of >30M mice, suggesting that decreased expression of C6 sulfating enzymes such as *CHST3* might be an important global mechanism behind the increase in C4S/C6S ratio. Interestingly, we observed an increase in the expression of *CHST13*, a carbohydrate sulfotransferase that is responsible for C4-sulfation of GAGs in the cortex but not hippocampus of 24-26M mice. This data suggests the brain area-specific regulation and provides an additional mechanism for the increase in the ratio of C4S/C6S in the cortex. Furthermore, we investigated the epigenetic changes in the regulation of the *CHST3* gene in a cell type-specific manner. We found that *CHST3* mRNA is significantly downregulated specifically in NeuN− cells that mostly represent the glial cell population, but not in neurons. These changes are accompanied by the decreased abundance of H3K4me3 on the promoter region of *CHST3* explaining the potential decrease in the expression of the gene.

## Acknowledgments

We thank Katrin Boehm and Jenny Schneeberg for technical support. This work has been supported by DAAD (fellowship to D.B.A), DFG (GRK 2413 “SynAge” to A.D.; Priority program 1738 to A.F.), the BMBF projects ENERGI (01GQ1421A to A.F. and A.D.), funds from the German Center for Neurodegenerative Diseases (to A.F. and A.D.). AF was supported by the ERC consolidator grant DEPICODE (648898); funds from the Hans and Ilse Breuer Foundation, and the Kleekamm Foundation of the University Medical Center Göttingen.

## Author’s contribution

D.B.A performed all QPCR experiments without cell sorting and did behavior experiment together with S.J. R.K and A.D designed the study and supervised data analysis. R.K wrote the manuscript and all authors contributed to the manuscript editing. A.F and M.S.S contributed with the analysis of cell type-specific epigenetic mechanism regulating the *CHST3* gene.

---

**Table 1.**
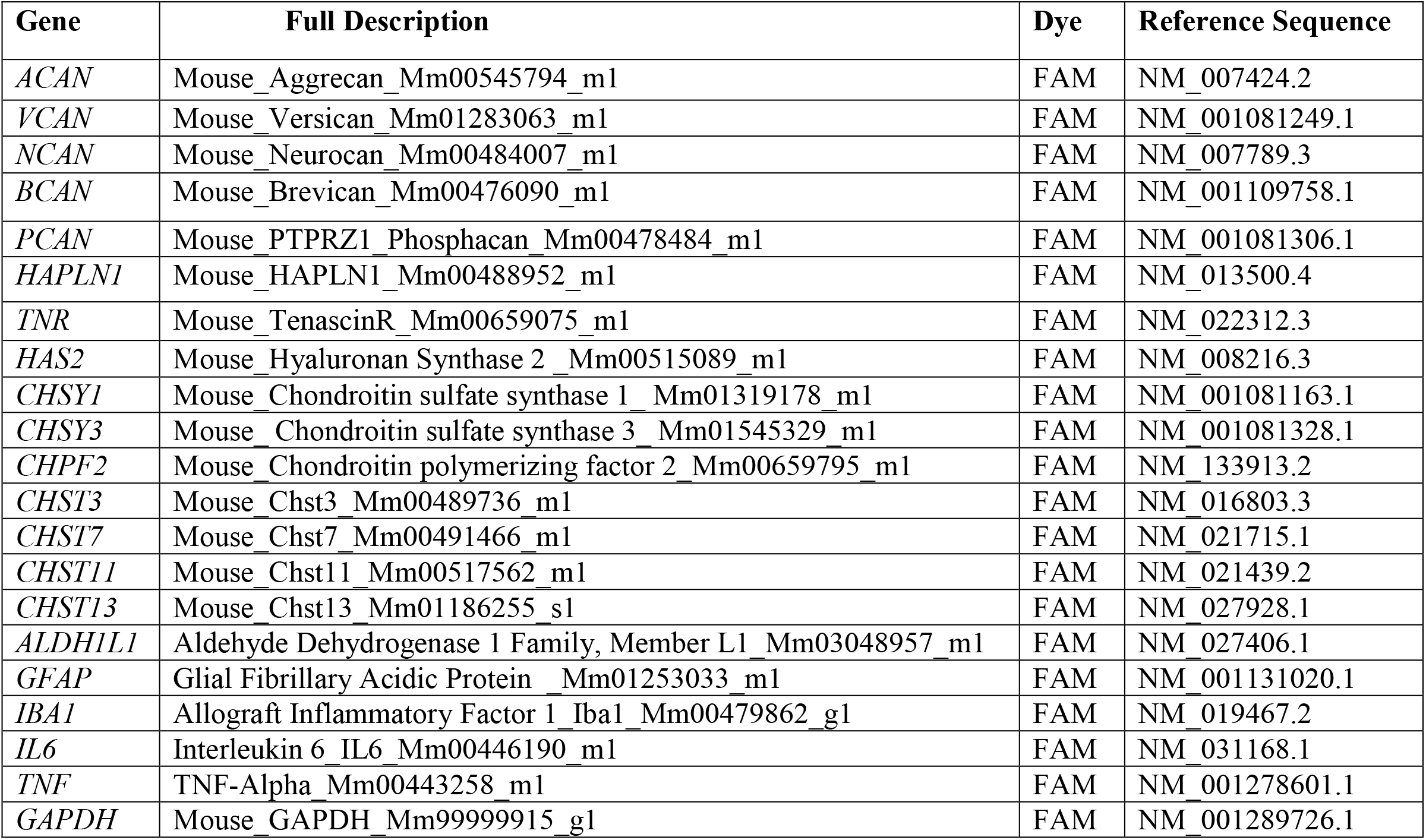
Taqman probes used for real‐time PCR analysis.

